# A microRNA cluster downstream of the selector gene Fezf2 coordinates fate specification with dendritic branching in cortical neurons

**DOI:** 10.1101/2021.11.03.467176

**Authors:** Asha Iyer, Verl B Siththanandan, Victoria Lu, Ramesh Nair, Lee O. Vaasjo, Maria J Galazo, Suzanne Tharin

**Affiliations:** Department of Neurosurgery, Stanford University, Stanford, CA, 94305 USA; Current address; Stanford Center for Genomics and Personalized Medicine, Stanford, CA, 94305, USA; Neuroscience program, Tulane Brain Institute, Tulane University, New Orleans, LA, 70118 USA; Department of Cell and Molecular Biology, Tulane University, New Orleans, LA, 70118 USA; Division of Neurosurgery, Palo Alto Veterans Affairs Health Care System, Palo Alto, CA, 94304 USA

## Abstract

In the cerebral cortex, cortical projection neurons comprise classes of neurons project to distant regions of the central nervous system. These neurons develop from the same progenitor pool, but they acquire strikingly different inputs and outputs to underpin strikingly different functions. The question of how corticospinal projection neurons - involved in motor function and implicated in paralysis - and callosal projection neurons - involved in cognitive function and implicated in autism - develop represents a fundamental and clinically important question in neurodevelopment. A network of transcription factors, including the selector gene Fezf2, is central to specifying cortical projection neuron fates. Gene regulation up- and down-stream of these transcription factors, however, is not well understood, particularly as it relates to the development of the major inputs to cortical projection neurons. Here we show that the miR-193b~365 microRNA cluster downstream of Fezf2 cooperatively represses the signaling molecule Mapk8, and impacts dendritic branching of cortical projection neurons.

## Introduction

The six-layered mammalian neocortex is generated in an inside-out fashion, with the earliest-born neurons populating the deepest layers, and the last-born neurons populating the most superficial layers (1). Corticospinal, interhemispheric callosal, and other cortical projection neurons develop through waves of neurogenesis, radial migration, and differentiation from radial glial progenitors of the ventricular zone and intermediate progenitor cells of the subventricular zone (2–5). In the mouse, corticospinal and a subset of callosal projection neurons are generated around embryonic day 13.5 (e13.5), and both reside in the deep cortical layer V. A larger subset of callosal projection neurons is generated around e15.5, and it populates the superficial cortical layer(s) II/III. A body of research from the last ~15 years has uncovered a set of transcriptional regulators that control cortical projection neuron development. These discoveries have been central to defining the current paradigm of combinatorial transcription factor controls (6–8).

FEZ Family Zinc Finger 2 (Fezf2) is a transcriptional regulator and selector gene required for specification of corticospinal and other deep-layer sub-cerebral projection neuron subtypes (9–15). Fezf2 regulates expression of the Ephrin type-B receptor 1 (EphB1) axon guidance receptor and the vesicular glutamate transporter 1 (vglut1) in corticospinal projection neurons (14). At the time of its discovery as a gene required for the subcortical projections of deep-layer cortical neurons, Fezf2 was also shown to be required for their dendritic development (9). Since that discovery, however, despite powerful and fruitful analyses of Fezf2 transcriptional targets (14), the molecular mechanism whereby Fezf2 controls dendritic development has remained elusive.

Here, we provide evidence for the role of a miRNA cluster downstream of the selector gene Fezf2 that serves as a potential link between neuronal fate specification and circuit inputs by regulating signaling pathways in dendritic branching. miRNAs are small noncoding RNAs that cooperatively repress multiple specific target genes post-transcriptionally (16). miRNAs regulate molecular programs that refine cellular identity. We have previously shown that miRNAs are selectively expressed in corticospinal vs. callosal projection neurons during their development, and that they can control cortical projection neuron fate (17). Here, we show that the miR-193b~365 miRNA cluster is differentially expressed by developing corticospinal vs. callosal projection neurons, that it is downstream of the transcription factor and selector gene Fezf2, that the cluster represses the callosal-expressed signaling gene Mitogen-Activated Protein Kinase 8 (MapK8), and that gain-of-function of the miR-193b and −365 miRNAs controls dendritic branching in cortical projection neurons in ways that phenocopy loss-of-function of Mapk8. To our knowledge, this is the first example of miRNA repression of a specific signaling pathway shaping neuron dendritic development and thereby circuit identity.

## Results

### The miR-193b~365 cluster is enriched in corticospinal projection neurons and is downstream of the selector gene Fezf2

We have previously shown that miRNAs are differentially expressed by corticospinal vs. callosal projection neurons during their development (17). When we examined the genomic organization of validated corticospinal-enriched miRNAs, we discovered that they mapped to two miRNA clusters: one on mouse chromosome 12, the 12qF1 miRNA cluster (17), and one on mouse chromosome 16, the miR-193b~365 cluster. We purified independent samples of corticospinal and callosal projection neurons in biological triplicate on postnatal day two (P2), and compared the Relative Quantity (RQ) of miRNA in corticospinal relative to callosal projection neurons, calculated as 2^−ΔΔCt^. The 2^−ΔΔCt^ method is particularly well suited to studying limiting quantities of input RNA, as is the case in pure populations of cortical projection neuron subtypes (18). We determined the RQ of each miRNA in corticospinal relative to callosal projection neurons using the small nucleolar RNA sno202 as a control. We found that miR-193b is 5,500-fold enriched and miR-365 is 2-fold enriched in corticospinal vs. callosal projection neurons on P2 (Figure 1A, D). It is not clear whether the observed orders of magnitude greater enrichment of miR-193b, vs. miR-365, in corticospinal projection neurons reflects differences in termination of transcription, in processing of a polycistronic transcript, or in miRNA degradation. To begin to understand how expression of the miR-193b~365 cluster is regulated, we carried out ATAC-Seq in purified corticospinal vs. callosal projection neurons on P2. Whereas the putative promoter region of the miR-193b~365 cluster is accessible in callosal projection neurons, it is inaccessible in corticospinal projection neurons. When this region is compared to the previously published results of Fezf2 ChIP-seq (14), it appears as though this region of inaccessibility in corticospinal projection neurons represents a Fezf2 footprint, strongly suggesting that selective expression of the the miR-193b~365 cluster by corticospinal, but not callosal, projection neurons is controlled by Fezf2.

**Figure 1.**
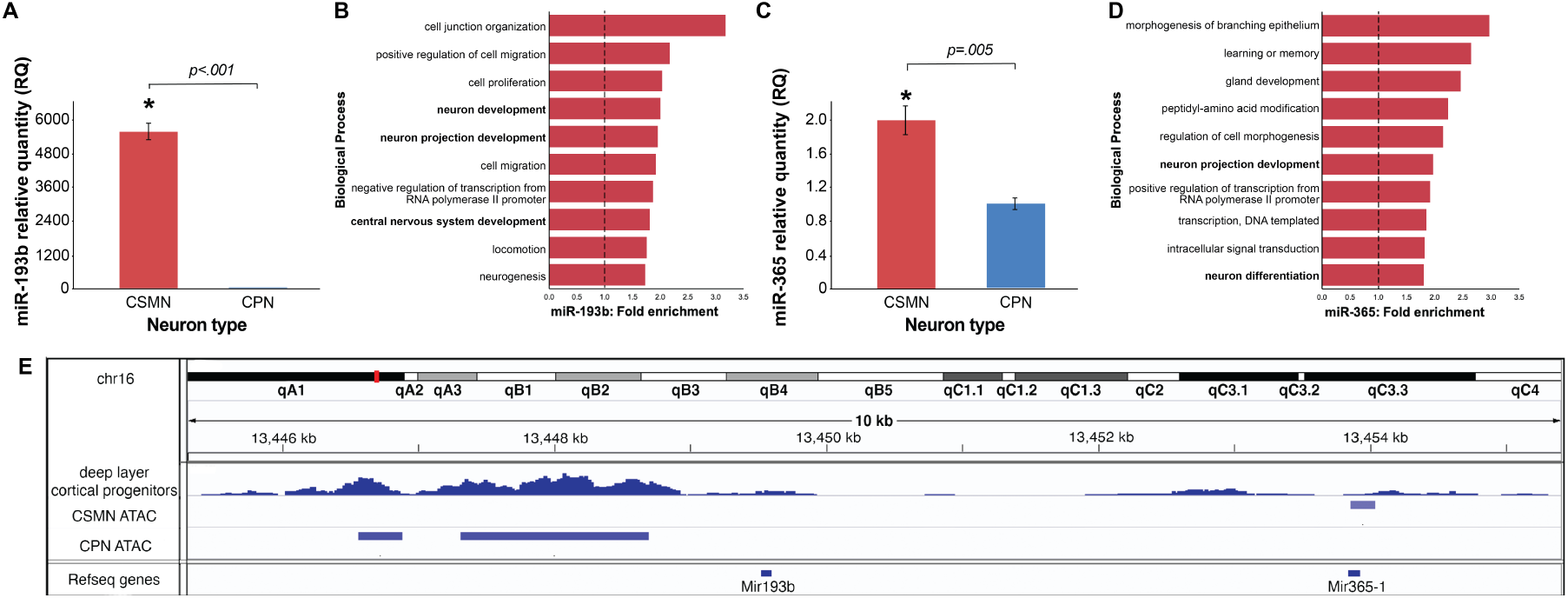
The miR-193b~365 cluster is enriched in corticospinal projection neurons and is downstream of the selector gene Fezf2. (A) miR193b-3p is 5630-fold enriched in Corticospinal projection neurons (CSMN) vs. Callosal projection neurons (CPN) on P2. (B) Top 15 over-represented biological processes that are predicted targets of miR-193b-3p by Gene Ontology (GO) analysis. (C) miR-365 is 2-fold enriched in CSMN vs. CPN on P2. (D) Top 15 over-represented biological processes that are predicted targets of miR-365 by Gene Ontology (GO) analysis. (E) Deep layer cortical progenitor Fezf2 ChIP-seq (14) combined with ATAC-Seq of purified CSMN and CPN reveal a Fezf2 footprint in the promoter region of miR-193b~365 cluster in corticospinal but not callosal projection neurons. Error bars represent SEM. * p<0.05

### miR-193b and miR-365 are predicted to act in neuron projection development and repress MapK8, a signaling molecule required for dendritic development

We carried out bioinformatic analyses to identify predicted targets of miR-193b and miR-365 using the search tools miRanda (19–22), Targetscan (23–27), DIANALAB (28–30), and miRDB (31, 32). Gene Ontology (GO) analysis of these predicted targets shows that neural development and neural process development are among the top ten overrepresented biological processes for both miR-193b and miR-365 (Figure 1C, D). Our bioinformatic analyses also specifically and consistently predicted that both miR-193b and miR-365 target the signaling molecule MapK8 (Figure 2A, D). MapK8 is important in neural development (33) and is preferentially expressed by callosal vs. corticospinal projection neurons (34), as would be predicted if miR-193b and miR-365 were repressing its expression in corticospinal projection neurons. To investigate whether miR-193b and miR-365 can repress MapK8 expression via the predicted sites in the 3’ UTR, we performed luciferase reporter gene assays in COS7 cells, as previously described (17, 35, 36). We used MapK8 reporter vectors containing either wild-type or mutated (mismatch) target sites and their flanking 3’UTR sequences. We found that miR-193b oligonucleotides significantly repress MapK8 luciferase reporter gene expression with wild-type, but not mismatch, miR-193b target sequences (Figure 2B). While we found that miR-365 oligonucleotides also repress MapK8 luciferase reporter gene expression with wild-type, but not mismatch, miR-365 target sequences, this result was not significant (Figure 2E), most likely due in part to low overall levels of reporter expression. Scrambled control miRNA oligonucleotides do not repress the MapK8 luciferase reporter gene.

**Figure 2.**
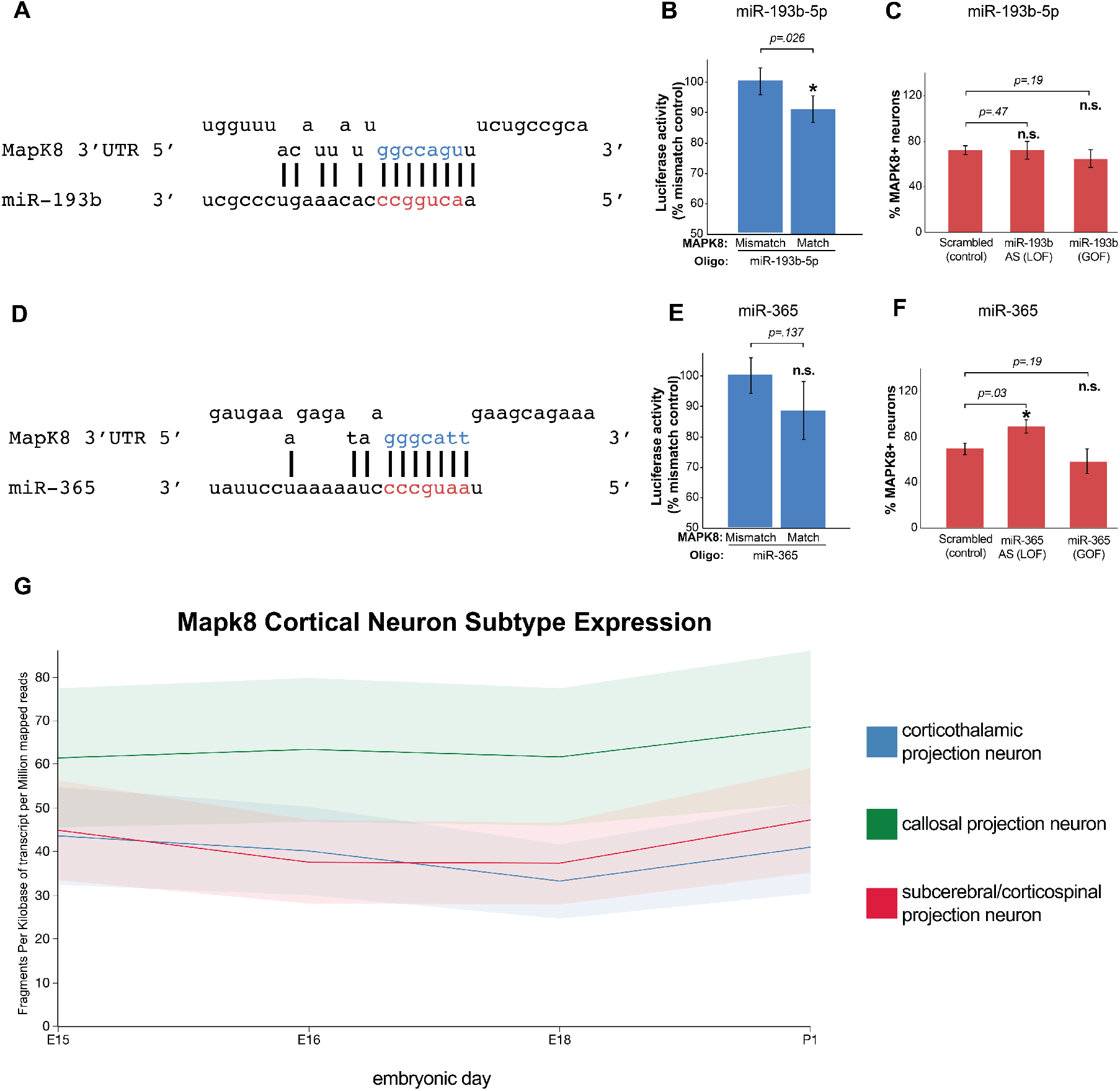
miR-193b and miR-365 repress MapK8. (A, D) Sequence alignments demonstrate that miR-193b and miR-365 are both predicted to target the MapK8 3’ UTR. Seed sequence base-pairing is shown in red. (B, E) miR-193b oligonucleotides repress a MapK8 3’UTR luciferase reporter gene bearing wild-type, but not mismatch, target sequences (B). miR-365 oligonucleotides do not significantly repress a MapK8 3’UTR luciferase reporter gene bearing wild-type or mis-match target sequences (D). Error bars represent SEM. * p<0.05 compared to mismatch control. n.s. not statistically significant compared to mismatch control. (C, F) miR-193b GOF and LOF show no change in the percent MapK8+ neurons compared to scrambled controls in embryonic cortical cultures (C). miR-365 LOF increases the percent MapK8+ neurons compared to scrambled controls in embryonic cortical cultures (F).

To test whether this finding extends to endogenous MapK8 in mouse cortical neurons, we performed gain-of-function (GOF) overexpression and antisense loss-of-function (LOF) experiments in cortical cultures. We used lentiviral vectors expressing miR-193b and GFP, miR-365 and GFP, antisense miR-193b and GFP, antisense miR-365 and GFP, or a scrambled miRNA and GFP. Cultures of e14 cortical cells were transfected with these vectors, as described in Materials and Methods, and were examined for MapK8 expression by immunofluorescence labeling on day 7 in culture, a stage considered to be equivalent to Postnatal day 1 (P1) *in vivo*. Double immunofluorescence with antibodies to MapK8 and GFP reveals that miR-365 LOF leads to an increase in the number of MapK8+ neurons in the targeted embryonic cortical cultures compared to control (Figure 2F), as would be expected if endogenous miR-365 were repressing Mapk8 in these cells. miR-193b GOF and LOF have no statistically significant effect on the number of MapK8+ neurons in the targeted embryonic cortical cultures compared to control (Figure 2C), although our luciferase reporter gene findings indicate an interaction of miR-193b and the MapK8 3’ UTR occurs (Figure 2B). Conversely, our luciferase reporter gene findings for miR-365 did not reach significance (Figure 2E), but support the trend seen in the miR-365 immunofluorescence studies. Collectively, the data indicate that miR-193b and miR-365 likely incrementally repress expression of the callosal-expressed signaling molecule MapK8 in cortical neurons, potentially regulating signaling during cortical projection neuron dendritic development in a subtype-specific way.

### miR-193b and miR-365 promote branching of neuronal projections in deep layer-enriched cultures

To better understand the roles of miR-193b and miR-365 in cortical neuron projection outgrowth and branching, we carried out overexpression GOF and antisense LOF experiments in cultured embryonic cortical neurons. Lentiviral vectors were used as described above. Cultures of e14 cortical cells were transfected with these vectors as described in Materials and Methods, and were examined for process morphology on day 7 in culture. To quantify the complexity of neural processes in transfected neurons, we performed a Scholl analysis. As described in Materials and Methods, we drew concentric circles around each neuron cell body examined and counted the number of intersections of branches at each circle (Figure 3A). GOF of both miR-193b and miR-365 result in a significant and substantial increase in the mean sum of intersections (expressed as a percent of control) compared to LOF (Figure 3B, C). Our results indicate that miR-193b and miR-365 promote branching of neuronal projections in primary cultures enriched for deep layer cortical neurons. In the motor cortex, MapK8 loss-of-function results in increased dendritic branching of layer V (predominantly corticospinal) projection neurons (37).

**Figure 3.**
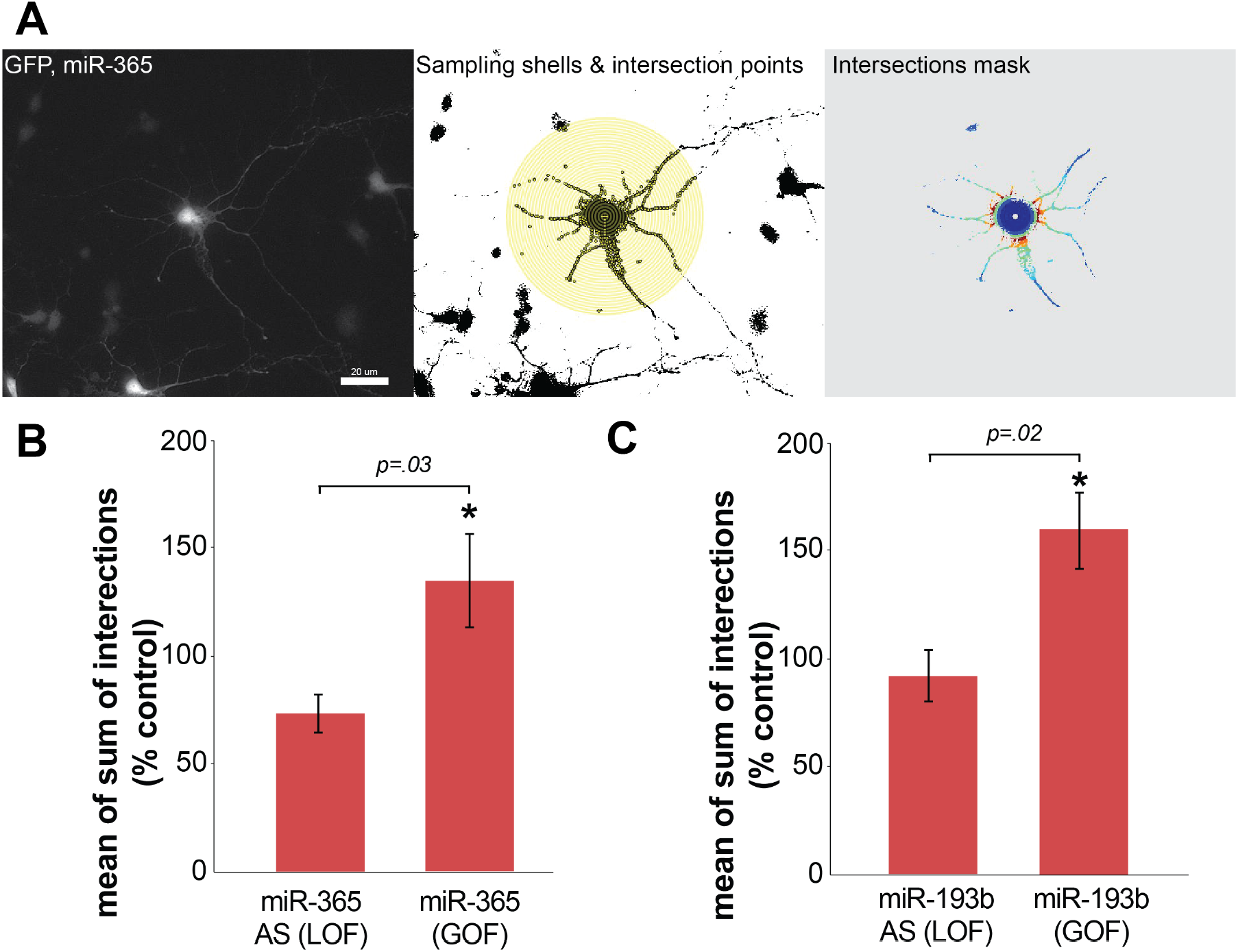
miR-193b and miR-365 promote branching of neuronal projections in cultures from deep layer progenitors. (A) Representative fluorescence micrograph of embryonic cortical cultures illustrates an increase in branching of neural processes with miR-193b GOF. Also shown, graphical outputs from Scholl plug-in on FIJI/IMAGEJ. Scale bar, 20 μm. (B) miR-365 overexpression gain-of-function (GOF) increases the percent the mean sum of intersections, and miR-365 antisense loss-of-function (LOF) decreases the mean sum of intersections, as a percentage of scrambled control in embryonic cortical cultures by Scholl analysis. (C) miR-193b overexpression gain-of-function (GOF) increases the percent the mean sum of intersections, and miR-193b antisense loss-of-function (LOF) decreases the mean sum of intersections, as a percentage of scrambled control in embryonic cortical cultures by Scholl analysis.

### Ectopic expression/gain-of-function of miR-365 in callosal projection neurons phenocopies MapK8 loss-of-function

To better understand the role of the miR-193b~365 miRNA cluster in shaping cortical dendritic development, we carried out miR-365 ectopic expression/gain-of-function experiments *in vivo* via *in utero* electroporation. We ectopically expressed miR-365, which is normally expressed in corticospinal but not callosal projection neurons, specifically in layer II/III callosal projection neurons by carrying out *in utero* electroporation at e14.5-e15, just prior to the peak of layer II/III callosal projection neuron production. Plasmid vectors over-expressing miR-365 and GFP, and similar vectors expressing a scrambled miRNA insert and GFP, were used. Constructs were injected into the embryonic lateral ventricle on e14.5-e15 for optimal incorporation into layer II/III callosal projection neurons. *In utero* electroporation was carried out as described in Materials and Methods. On P0, embryos were removed, fixed, sectioned, and examined by immunocytochemistry for dendritic branching. We found that ectopic expression of miR-365 in callosal projection neurons results in decreased dendritic branching compared to control (Fig 4 A-E). *In vivo*, MapK8 loss-of-function results in decreased dendritic branching of layer II/III (predominantly callosal) projection neurons, compared to wild-type (37). Our observed miR-365 ectopic expression/gain-of-function dendritic phenotype therefore represents a phenocopy of the previously described MapK8 loss-of-function dendritic phenotype during development (37), as would be expected if an important role of miR-365 in cortical projection neurons is repressing MapK8.

**Figure 4.**
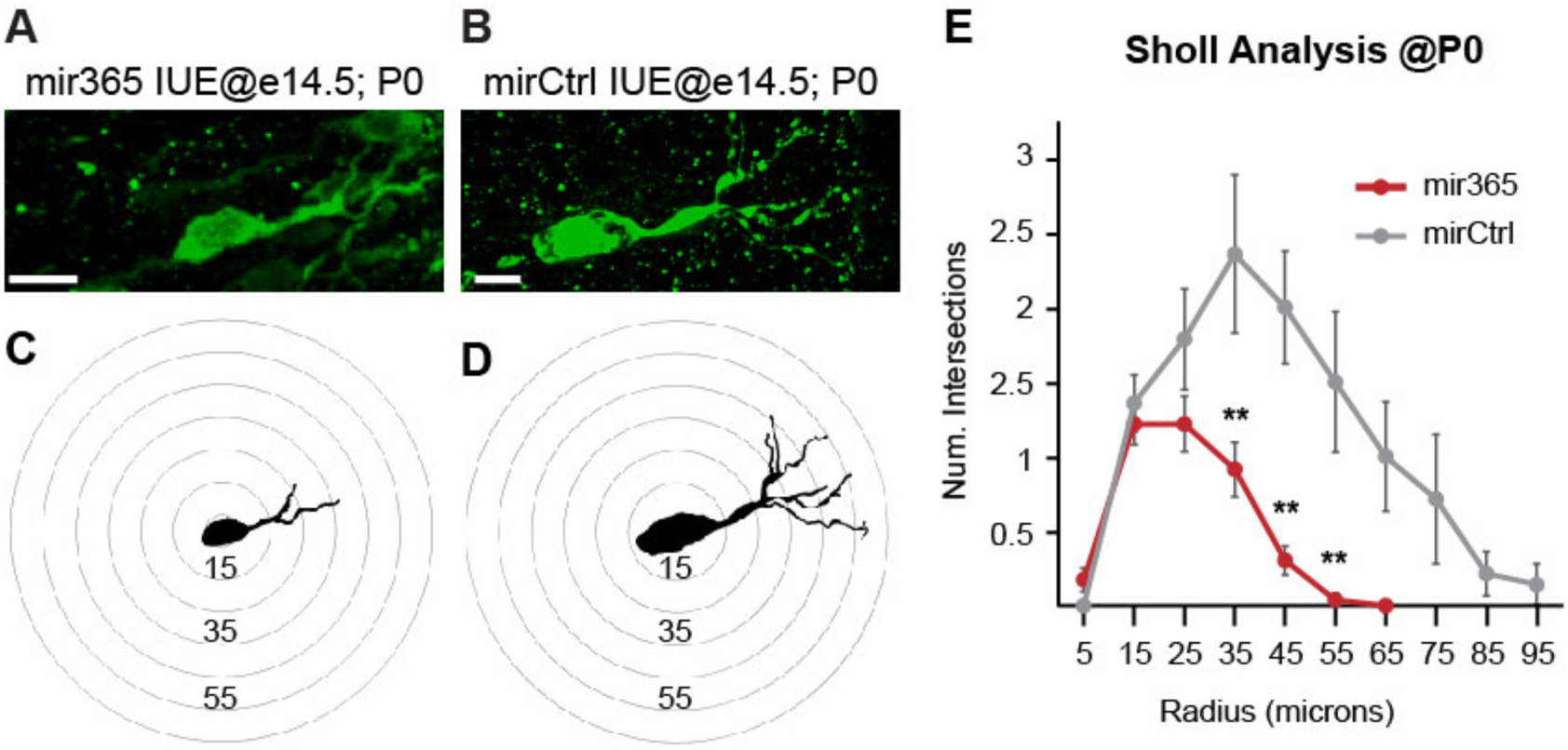
Ectopic expression of miR-365 in callosal projection neurons phenocopies Mapk8 loss of function. Representative fluorescence micrographs of P0 cortices show decreased dendritic branching in miR-365 GOF (A) compared to control (B). Scale bar, 10μm. Scholl analysis of the neurons pictured in A,B again shows decreased dendritic branching in miR-365 GOF (C) compared to control (D). Numbers represent distance from the soma in μm. Quantitative Scholl analysis at P0 shows a significantly reduced number of intersections in miR-365 GOF compared to control, beginning at a distance of 35μm from the soma (E). Error bars represent SEM. t-test **p<0.05 compared to scrambled control. Reconstructed neurons, N=23 for miR365, N=14 control.

## Discussion

Corticospinal and deep-layer callosal projection neurons arise from the same progenitor pool at the same time, yet develop highly divergent inputs and outputs to serve highly divergent functions. While a set of canonical transcriptional regulators, notably including Fezf2, SATB2, and CTIP2, has been shown to control corticospinal and callosal projection neuron development, the evidence linking these transcriptional regulators to corticospinal and callosal dendritic development is lacking. Here we provide the first evidence that cooperative refinement of signaling mechanisms by clustered miRNAs regulates dendritic development in cortical projection neurons. We identify that the miR-193b/365 cluster, previously shown to be enriched in corticospinal projection neurons during their development (17), (1) is downstream of the selector gene Fezf2, (2) can repress the signaling molecule Mapk8, and (3) can control branching of cortical dendrites according to, and shaping, cortical projection neuron subtype.

MapK8 (JNK1) is a member of a subfamily of mitogen activated protein kinases (MAP kinases) that are involved in intracellular signaling and highly expressed in the nervous system (38). In the motor cortex, MapK8 loss-of-function results in increased dendritic branching of layer V pyramidal neurons (predominantly corticospinal projection neurons) and decreased dendritic branching of layer II/III pyramidal neurons (primarily callosal projection neurons), compared to wild-type (37). Consistent with these findings, we observe in culture that gain of function of miRs-193b and −365 - expected to phenocopy loss of function of their shared target MapK8 - result in increased process branching of pyramidal neurons transfected within the peak of corticospinal projection neuron production but prior to the major peak of callosal projection neuron production (Figure 3). While these cultures are expected to be enriched for deep-layer neurons, it is possible that our observed results represent an underestimate of the effect of miRs-193b and −365 loss and gain of function on corticospinal projection neurons, since there are likely superficial layer callosal projection neurons also present in these cultures.

MapK8 is also specifically required for netrin-1 attractive axon guidance signaling through its receptors DCC and DSCAM in the developing brain (39). Netrin-1 signaling via DCC is required during callosal projection guidance (40). Indeed, Tang and Kalil had previously shown decreased axonal branching in the setting of MapK8 inhibition in cultured neonatal, cortical neurons (41). Future experiments, including examination of axonal branching, will be important to determine whether the corticospinal-expressed miR-193b and miR-365 cooperatively repress this callosal-expressed netrin-1 attractive midline crossing axon guidance pathway in developing corticospinal projection neurons, via MapK8 repression.

Clustered miRNAs often cooperatively repress the same gene and/or interacting genes within a pathway (42). Both in luciferase assays and in immunocytochemical analyses of primary cortical cultures, we observe partial repression of MapK8 by the miR-193b/365 cluster. Curiously, repression of the Mapk8 reporter in luciferase assays reached statistical significance for miR-193b but not miR-365, whereas repression of the Mapk8 protein in primary cortical cultures reached statistical significance for miR-365 but not miR-193b. This observation, combined with the modest effect sizes, suggest that optimal repression of Mapk8 by the miR-193b/365 cluster is co-operative and may require the activity of both miR-193b and miR-365, consistent with known cooperative repression mechanisms of miRNA action (42, 43).

The discovery that Fezf2 is required for normal corticospinal projection neuron dendritic morphology provided evidence that this selector gene controlled all aspects of deep layer projection neuron development (9). However, since that time, the connection between early dendritic development and Fezf2, or any of the other canonical transcription factors of cortical projection neuron development, remains unclear. Here we show that the miR-193b/365 microRNA cluster downstream of Fezf2 coordinates fate specification with dendritic branching in cortical neurons, likely in part via coordinate repression of the signaling molecule Mapk8. In support of this model, miR-193b and −365 loss-of-function results in reduced branching of predominantly deep-layer cortical projection neurons in primary culture, a phenocopy of the deep-layer cortical projection neuron dendritic phenotype of Fezf2 loss-of-function *in vivo* (11). This is expected if the miR-193b~365 cluster is downstream of Fezf1 in cortical projection neuron dendritic development. miR-193b and −365 gain of function, on the other hand, result in increased branching of deep-layer cortical projection neurons in primary culture, a phenocopy of the deep-layer projection neuron dendritic phenotype of Mapk8 loss-of-function during dendritic development *in vivo* (37). This is expected if the miR-193b~365 cluster is upstream of Mapk8 in cortical projection neuron dendritic development. *In vivo*, ectopic expression/gain-of-function of miR-365 in callosal projection neurons phenocopies Mapk8 loss-of-function in these neurons, again as predicted by our model in which the miR-193b~365 cluster is upstream of, and represses, Mapk8. Taken together, our results suggest a role of subtype-specific miRNAs in coordinating fate specification with circuit identity, in this case by shaping projection neuron inputs during cortical development.

## Acknowledgements

We thank members of the ST and MJG labs for scientific discussions and helpful suggestions. This work was supported by grants from the NIH (K08 NS091531), AOSpine North America (Young Investigator Research Grant Award), a Stanford McCormick Faculty Award, and a Stanford Maternal and Child Health Research Institute Pilot Award, and a Stanford Maternal and Child Health Bridge Funds award to ST. LOV is supported by Louisiana Board of Regents funds (BoR RCS program) to MJG. MJG is supported by grants from the Brain and Behavior Foundation (NARSAD young investigator award), and Carol Lavin Bernick Faculty Grant Program. ST is a Tashia and John Morgridge Endowed Faculty Scholar in Pediatric Translational Medicine of the Stanford Maternal and Child Health Research Institute. We acknowledge the Stanford Neuroscience Microscopy Service, supported by NIH NS069375.

## Author Contributions

ST, and AI conceived the project; ST, AI, VL, MJG, and VBS designed the experiments; VBS, ST, MJG, LOV, and AI performed the experiments; VBS, ST, MJG, and AI analyzed and interpreted the data; AI performed the bioinformatic analyses; RN and ST performed the ATAC-Seq analysis; ST, MJG, and VBS made the figures; ST, AI, MJG, and VBS synthesized and integrated the findings and wrote and revised the paper.

## Declaration of Interests

The authors declare no competing interests.

## Materials and Methods

### CSMN and CPN purification

All animal work was carried out in compliance with Stanford University Institutional Animal Care and Use Committee approved protocols and institutional and federal guidelines. CSMN and CPN were purified from C57BL/6J mice as previously described (10, 44–46). Briefly, CSMN were retrogradely labeled at P0 or P1 from the cerebral peduncle by injection of cholera toxin B subunit (CTB) from *Vibrio cholerae* with FITC conjugate (List Biological Laboratories, Campbell, CA) under ultrasound microscopic guidance. At P1 or P2, neuron bodies in the motor cortex were isolated by microdissection of deep cortical layers, and dissociated to a single cell suspension by papain digestion and mechanical trituration. CPN were retrogradely labeled on P0 or P1 by injection of CTB into the contralateral hemisphere under ultrasound microscopic guidance. On P1 or P2, labeled cortex was microdissected and dissociated to a single cell suspension by papain digestion and mechanical trituration. FACS purifications were performed by the Stanford Shared FACS Facility. For differential miRNA expression analysis, neurons in suspension were FACS-purified into RNAlater (Life Technologies) using a FACSVantage sorter (BD), and purified labeled neuron cell bodies were then frozen at −80°C until RNA purification. For ATAC-seq library preparation, the neuron suspensions were sorted into cold 50/50 media (50% DMEM, Life Tech 10569-010 and 50% Neural Basal Media, Life Tech 21103-049) for immediate processing.

### RNA purification

microRNA was extracted from purified projection neuron cell bodies using the Ambion miRVana microRNA isolation kit (Ambion, Austin, TX) according to the manufacturer’s instructions. RNA quality was analyzed on a BioAnalyzer 2100 (Agilent), and confirmed to be excellent.

### Differential miRNA expression analysis

miRNAs were expression was analyzed via qPCR from purified P2 CSMN and CPN using sno202 as a control. Relative Quantity (RQ) was calculated for each of the miRNAs, with RQ = 2^−ΔΔCt^, providing a measure of the fold difference in miRNA expression by one cell type vs. the other. We performed experiments in biological triplicate.

### ATAC-seq Library Preparation and Sequencing

Preparation of ATAC-seq libraries from viable cells has previously been described(47). Briefly, viable FACS sorted cells, 50,000 per biological replicate, were lysed in non-ionic detergent to isolate the nuclei. The nuclei were suspended in a transposition mix (TD Tagment DNA Buffer, Illumina Catalog No. 15027866, and TDE1 Tagment DNA Enzyme, Illumina Catalog No. 15027865), and incubated at 37°C to generate DNA fragments. After purification (Qiagen MinElute PCR purification Kit) the DNA fragments could be stored at −20°C. The DNA fragments were amplified using indexing primers, where the required number of PCR cycles is determined by qPCR (the required cycles was 25% of maximum fluorescent intensity). The product was purified once more to give the final library. Library concentrations were in the nano-molar range. The library quality was assessed using a BioAnalyzer (2100 Agilent, Stanford Functional Genomics Core). Samples with an RNA integrity number above 9 were used for ATAC-seq library sequencing. ATAC-seq libraries were sequenced by the Stanford Functional Genomics Core on an Illumina MiSeq Sequencer. Each library produced approximately 200M, 150-bp paired-end reads.

### ATAC-seq Analysis

The ENCODE ATAC-seq uniform processing pipeline v1.4.2 was used for replicated and unreplicated data (*https://www.encodeproject.org/atac-seq/*). The singularity-based pipeline for automated end-to-end quality control and processing of ATAC-seq data was installed from GitHub (*https://github.com/ENCODE-DCC/atac-seq-pipeline*). Alignments of paired-end fastq reads were performed against the mouse mm10 reference. A statistical procedure called the Irreproducible Discovery Rate (IDR), which operates on the replicated peak set and compares consistency of ranks of these peaks in individual replicate/pseudoreplicate peak sets, was used to generate a higher confidence, reproducible set of peaks with low false positive rates (48). IDR can operate on peaks across a pair of true replicates resulting in a “conservative” output peak set, or across a pair of pseudoreplicates resulting in an “optimal” output peak set. These IDR conservative/optimal peaks for true/pseudo replicates respectively were then viewed on the Integrative Genomics Viewer (IGV) (49).

### miRNA target prediction and Gene Ontology (GO) analysis

We used miRanda (19–22), Targetscan (23–27), DIANALAB (28–30), and miRDB (31, 32) to search for predicted miRNA targets. GO analysis was performed using the Panther GO tool (50).

### Luciferase assays

Luciferase reporter assays were performed using the Dual-Glo Luciferase Assay System (Promega), pmir-GLO based reporter constructs, and microRNA oligonucleotides (Horizon Discovery) according to the manufacturer’s instructions, as previously described (17, 35, 36). Briefly, COS7 cells (10^4^/well) were seeded in a white 96-well plate. The following day, pmir-GLO reporter-miRNA oligo-DharmaFECT Duo (Dharmacon) transfection mixtures were prepared. The media from the 96-well plate was replaced with the transfection mixture, and the plate was incubated overnight. Firefly and renilla luciferase reporter fluorescence was read using a Tecan Infinite M1000 (Stanford High-Throughput Bioscience Center Core Facility). The ratio of firefly to renilla fluorescence was calculated for each well. Averages were compared for triplicates of each condition. Match reporter vectors contained the two wild-type predicted miR-193b (TACATTATTGGCCAGTTTCTGCCGCA) or miR-365 (ATAGGGCATTGAAGCAGAAA) seed regions with 30bp of flanking MapK8 3’ UTR on either side of each. Mismatch reporter vectors were identical to match reporters except that the seed sequences were replaced by GGGGGGG. The experiments were replicated in n=5 independent cultures.

### Lentivirus vectors

Lentivirus vectors were modified from the pSicoR backbone (51), a gift from Tyler Jacks (Addgene plasmid # 11579). Expression of miRNA was under direction of the strong U6 promoter. miRNA inserts were: miR-193b (gain of function: AACTGGCCCTCAAAGTCCCGCTTTTTT), antisense miR-193b (loss of function: AGCGGGACTTTGTGGGCCAGTTTTTTT), miR-365 (gain of function: TAATGCCCCTAAAAATCCTTATTTTTT), antisense miR-365 (loss of function: ATAAGGATTTTTAGGGGCATTATTTTT), or scrambled (control, CCTAAGGTTAAGTCGCCCTCGCTCCGAGGGCGACTTAACCTTAGGTTTTT). All miRNA inserts were cloned between Hpal and Xhol sites. Expression of GFP was under direction of the CMV promoter. Lentivirus packaging was provided by System Biosciences (Palo Alto, CA). Titers of VSV-G pseudotyped viral particles were ~10^7^ IFUS/mL.

### Cortical cultures

Embryonic cortical cultures were prepared as previously described (17, 52, 53). Briefly, e14 cortices were dissected and gently dissociated by papain digestion. A single cell suspension was prepared and plated onto poly-D-lysine (100μg/ml, Sigma) and laminin (20μg/ml, Life Technologies) coated coverslips in cortical culture medium. Cells were infected with lentivirus, and cultured on coverslips placed in 6-well plates for 7 days in growth media (50% DMEM, 50% neural basal media, supplemented with B27, BDNF, forskolin, insulin, transferrin, progesterone, putrescine, and sodium selenite). Under these conditions we observe >~95% neurons and very few (<~5%) astrocytes.

### Immunocytochemistry of cultured cells

Cells were fixed with 4% paraformaldehyde (PFA) in PBS. Coverslips were blocked with PBS containing 0.1% Triton-X100, 2% sheep serum, and 1% BSA, and cells were then incubated with primary antibodies: anti-MapK8 (Abcam, mouse monoclonal, dilution 1:200). Following wash steps, the cells were incubated with secondary antibodies conjugated with fluorophores: anti-mouse (Pierce, CY3-conjugate, dilution 1:1000), and anti-GFP (AbCam, goat-FITC conjugated, dilution 1:250). Cells were re-fixed in 4% PFA, and washed thoroughly in ddH2O. The coverslips were then mounted on microscope slides using Fluoroshield with DAPI (Sigma), and left overnight to dry. The following day, slides were sealed with clear nail polish, and imaged on a Zeiss AxioImager microscope.

### Scholl analysis of cultured neurons

We analyzed neurons in 16 randomly selected high-powered fields, blinded to experimental condition. The experiments were replicated in 5 independent cultures. Statistical analyses were carried out using FIJI/IMAGEJ on micrographs that were converted to greyscale, and image thresholds adjusted to highlight neural processes. Individual neurons – isolated from surrounding processes – were selected and the radius from the soma to the furthest clear process draw using the line segment tool. The Sholl plugin was used to draw concentric sampling shells along the length of the drawn line. The radius of the first shell was set to exclude the soma (~10 μm) and subsequent shell radii increased in 5 μm steps. The number of intersection points at each sampling shell were recorded and summed. Comparisons were made between the mean of the sums of intersections for hundreds of cells for LOF and GOF, expressed as a percentage of the control.

### *In utero* electroporation

For miR-365 gain-of-function experiments, a CAG/H1 promoter plasmid was used to drive expression of GFP and mature miR-365. For control experiments, a similar plasmid was used to drive expression of GFP and a scrambled control microRNA (scrambled control). *In utero* electroporation of embryonic mouse cortical neurons was carried out as previously described (10).

Briefly, 1 μL of plasmid DNA at 1 μg/μL mixed with 0.01 % Fast Green was injected into the lateral ventricle of e15 CD1 mouse embryos *in utero*. Plasmids were electroporated into the neocortical ventricular zone using 5mm diameter platinum disk electrodes and a square-wave electroporator (BTX ECM 830) with five 30-Volt pulses of 50 milliseconds duration at 950 millisecond intervals, as previously described (54). Electroporated embryos were collected for analysis at P0, and their brains fixed by immersion in 4% PFA overnight.

### Immunocytochemistry of brain sections

Embryonic brains were drop-fixed in 4%PFA and post-fixed overnight in 4%PFA at 4°C. Brains were embedded in 4% agarose and sectioned coronally at 75 μm on a vibratome (Leica VT1000S). Tissue sections were collected serially and stored in PBS with 0.0025% Sodium Azide (NaN3) at 4°C. Immunolabeling was performed to enhance GFP. Tissue sections were washed in PBS, and incubated for 1-2 hours at room temperature in a solution containing 1.5% Triton-X, 0.0025% NaN3, 0.3% bovine serum albumin (BSA), and 8% normal goat serum (NGS) in PBS for permeabilization and blocking non-specific binding of antibodies. Sections were incubated in primary antibody overnight at 4°C (rabbit anti-GFP 1:1000; Invitrogen A-6455 RRID:AB_221570), and in secondary antibody at 1:500 for 3 hours at room temperature goat (anti-rabbit Alexa Fluor 488, cat#A11008 Invitrogen). Sections were mounted using Vectashield with DAPI.

### Microscopy and image analysis

Images were acquired a Nikon C2 confocal microscope. Tile-scan epifluorescence images were acquired at 20x magnification (NA 0.75) to document the electroporated area. Only electroporated neurons in the somatosensory cortex were selected for further analyzed. Z-stack tile-scan images containing multiple electroporated neurons were acquired with a 60x (1.40) at 0.5 μm Z-resolution, 1024 × 1024 pixels lateral resolution, and 2-frame averaging. Deconvolution was performed (Richardson-Lucy Deconvolution algorithm, for a maximum of 30 iterations) with Nikon Elements. Tile images at 20x and 60x were stitched with 15% overlap using Optimal Path.

Neurons in the somatosensory cortex with sufficient GFP signal to detect second or higher order branching were selected for analysis.

Somas and dendrites were 3D-reconstructed using Neurolucida 360 (MBF Bioscience) and subjected to Sholl analysis using Neurolucida Explorer v.2021.1.1. (MBF Bioscience). Sholl analysis was performed for intersections using 10 μm concentric circles surrounding the cell soma. The number of intersections per radius increment were measured. Two tailed t-test was used to determine significant differences for the Sholl intersections. Imaging acquisition and neuron reconstruction were performed blinded to the experimental conditions.

## Notes

### Competing Interest Statement

The authors have declared no competing interest.

